# Modelling phage-bacteria interactions driving predation and horizontal gene transfer

**DOI:** 10.1101/291328

**Authors:** Jorge A. Moura de Sousa, Ahlam Alsaadi, Jakob Haaber, Hanne Ingmer, Eduardo P.C. Rocha

## Abstract

Bacteriophages shape microbial communities by predating on them and by accelerating their adaptation through horizontal gene transfer. The former is the basis of phage therapy, whereas the latter drives the evolution of numerous bacterial pathogens. We present a novel computational approach (eVIVALDI – eco-**eV**olutionary m**I**crobial indi**V**idu**AL**-base**D** s**I**mulations) to study phage-bacteria ecological interactions that integrates a large number of processes, including population dynamics, environmental structure, genome evolution, and phage-mediated horizontal transfer. We validate and illustrate the relevance of the model by focusing on three specific questions: the ecological interactions between bacteria and virulent phage during phage and antibiotic therapy, the role of prophages as competitive weapons, and how autotransduction facilitates bacterial acquisition of antibiotic resistance genes upon lysis of antibiotic resistant competitors. Our model recapitulates experimental and theoretical observations and provides novel insights. In particular, we find that environmental structure has a strong effect on community dynamics and evolutionary outcomes in all three case studies. Strong environmental structure, relative to well-mixed environments and especially if antibiotics are heterogeneously distributed, enhances the rate of acquisition of resistance to both phages and antibiotics, and leads to more accurate predictions of the dynamics of lysogen invasion in the gastrointestinal tract. We predicted the co-existence of invaders and resident lysogens in autotransduction under a range of parameters, and validated this key prediction experimentally. By linking ecological and evolutionary dynamics, our modelling approach sheds light on the factors that influence the dynamics of phage-bacteria interactions. It can also be expanded to put forward novel hypotheses, facilitating the design of phage therapy treatments and the assessment of the role of phages in the spread of antibiotic resistance.

**AUTHOR SUMMARY:** In the face of a growing threat of antibiotic resistant bacteria, bacteriophages have re-emerged as a potential alternative to clinical treatments of infections, as they are efficient bacterial predators. However, bacteriophages can also promote, through a mechanism called transduction, the dissemination of adaptive traits between bacteria, including antibiotic resistance genes. Importantly, these two types of interactions (predation and transduction) can co-occur, which creates difficulties in predicting their outcome. We have developed eVIVALDI (eco-**eV**olutionary m**I**crobial indi**V**idu**AL**-base**D** s**I**mulations), a computational model that allows the simulation of microbial communities with a focus on the mechanisms involved in phage-bacteria interactions, across time and in different types of environments. eVIVALDI can be used to understand the conditions where phages are more likely to be successfully used to eliminate bacteria or, in the other hand, the conditions where they increase the probability of dissemination of adaptive traits. Our research highlights the importance of considering the diverse ways that phage and bacteria interact, and the relevant ecological conditions where these interactions take place, to understand how bacteriophages shape microbial communities and how they can be used as a clinical tool.

## Introduction

Microbial organisms are pervasive across all natural environments, including the human body. Their adaptation and organization in communities may lead to disease [1], drive host evolution [2], and produce major changes in ecosystems [3,4]. Ecological interactions in microbial communities influence, and are influenced by, the rapid pace with which microbes acquire adaptive changes [5,6]. A striking example is the relationship between bacteria and bacteriophages (from here on referred to as phages), because the latter predate on the former whilst also driving their adaptation [4]. Phages are the most abundant entities in nature [7,8] and very efficient bacterial predators; it has been estimated that they promote the turnover of ~20% of bacterial mass every single day in certain environments [9,10]. In the context of widespread antibiotic resistance, this has led to a rekindled interest in phage therapy as an adjuvant or a replacement of antibiotic therapy against multi-resistant bacteria [11].

Virulent phages follow a strictly lytic cycle within their hosts, but they often exist in diverse communities with other virulent and temperate phages. Infection by the latter can lead to either the lytic cycle or their integration in bacterial genomes, as prophages (lysogenic cycle). Temperate phages are not used for phage therapy because lysogeny prevents them from extinguishing bacterial populations and confers resistance to closely, and sometimes distantly related phages – a mechanisms known as superinfection exclusion [12–14]. However, half of all bacterial genomes contain at least one, and up to 20, prophages, with these being more frequently found in bacterial pathogens [15], which means that they cannot be ignored in phage therapy. The expression of prophage genes may provide novel traits to the host (lysogenic conversion), and many cases have been described where prophages carry adaptive traits implicated in virulence or resistance to stress [16]. Virions arising from prophage induction can infect closely related competitor bacteria that are non-lysogenic for the phage, decreasing bacterial competition, increasing prophage frequency, and liberating resources that can be used for the growth of the remaining lysogenic population [17]. In this case, prophages have been regarded as weapons against bacterial competitors [13,18].

Phages can drive horizontal gene transfer between bacteria by transduction [19]. This can be a hazard in the case of phage therapy if the transferred traits are virulence factors or antibiotic resistance genes. Specialized transduction occurs in temperate phages when prophage excision leads to the transfer of neighboring chromosomal genes. Generalized transduction occurs when bacterial DNA is delivered to other cells after being encapsulated in virions, due to the specificities of the *pac* DNA packaging system [20]. Although these mechanisms are commonly used as genetic engineering tools [21], they have been considered inevitable consequences of errors in phage replication machineries and their rates in nature are poorly known. In the lab, they vary across several orders of magnitude (between 10^-11^ and 10^-3^ [22,23]), depending on the phage, the environment, and the type of culture media [19]. Importantly, phage driven transmission of bacterial DNA can have particularly nefarious consequences for humans. Transducing phages are responsible for the transmission of virulence factors in *Staphylococcus aureus* [24], and may accelerate the spread of antibiotic resistance genes [25,26]. Transduction can also have an impact at very short time scales: prophage induction facilitates the capture of adaptive traits (e.g., an antibiotic resistance gene) from a second bacterial strain that is infected by the phage and, through generalized transduction, transfers the gene back to the lysogenic strain. This process has been called autotransduction [27]. Hence, phages drive the evolution of bacterial gene repertoires and may spread virulence or antibiotic resistance factors during phage therapy.

The diversity of interactions between phages and bacteria may obscure the effects of each of them. Experimental approaches have described and clarified the mechanisms underlying these interactions, but usually focused on simplified environments [8,28]. *In vivo* studies of these interactions (e.g., in mammalian hosts [29]) tackle more natural environments, but have limited resolution in tracking temporal dynamics or the effects of individual mechanisms. Mathematical modelling provides a complementary approach to the study of phage-bacteria interactions, providing important insights on their co-evolutionary processes [30] or the dynamics of particular bacterial defense mechanisms [31,32]. Previous models have focused on individual mechanisms of interaction in simple environments (*e.g*., how the evolution of resistance to phage can affect clinical treatments [33]), because tackling multiple mechanisms and spatial heterogeneity hinders the development of analytical solutions. Yet, natural communities, and particularly those relevant for phage therapy, include complex interactions and spatial structure [34–38]. This may explain why models sometimes fail to fully reproduce *in vivo* dynamics of phage infection [28], and why there is paucity of models on the impact of phage-mediated horizontal gene transfer in the adaptation of bacterial communities (but see the work of Volkova *et al*. [39] for a theoretical comparison between the relative efficacy of transduction versus conjugation in transmitting an adaptive trait).

Individual-based models (IBMs) are an alternative to population-based mathematical approaches for studying complex microbial systems [40]. They have been useful to understand, for instance, the effect of spatial structure in microbial social evolution [41] or the interactions between bacteria and virulent phages in biofilms [36]. Although computationally intensive, IBMs provide a framework to study biological systems through the incorporation of different (and potentially interacting) mechanisms at the level of the individual. Population-level dynamics can then emerge from the collective individual behaviors. This makes IBMs particularly appealing to investigate phage-bacteria interactions, because these involve both ecological (e.g., predation) and evolutionary (e.g., transduction of adaptive traits) scales, with antagonistic mechanisms defined at the individual level (e.g., the lysis-lysogeny decision of temperate phage). To study these multiple roles of phages in microbial communities, we developed an IBM approach that is able to simulate diverse mechanisms and eco-evolutionary contexts: eVIVALDI – eco-**eV**olutionary m**I**crobial indi**V**idu**AL**-base**D** s**I**mulations. We focus on three questions, of gradually increasing complexity, that are relevant for bacterial evolution and phage therapy, and that cover a range of possible eco-evolutionary interactions between bacteria and phage. First, we introduce the basic scheme of the simulation with the study of ecological interactions between co-evolving virulent phages and bacteria under phage and antibiotic pressure in structured environments. Then we introduce lysogeny and super-infection exclusion in the model to study the role of prophages as competitive weapons. We show that our model provides better fit to previous experimental results than earlier models. Finally, we introduce transduction and the way we encode individual genomes in the model to elucidate how bacteria may obtain novel adaptive genes from sensitive bacteria by autotransduction. We use eVIVALDI to explore and quantify the different mechanisms of phage-bacteria interactions and to gain insights on how the structure of the environment can affect these interactions and the community dynamics. We tackle each question by demonstrating the ability of the model to capture previous results and then show how its complexity highlights new relevant features.

## Methods

### Concept and basic implementation

The eVIVALDI model was developed in Python (version 2.7.3), using an object-oriented approach, with a focus on the flexibility and extensibility of mechanisms and parameters simulated. The complete ODD (Overview, Design concepts, and Details) protocol [42] of the developed model is available as supplementary text (Text S1, Table S1 has the parameters modelled and their possible values), but below is a brief overview of the model. The source of the software can be obtained in the following link: https://gitlab.pasteur.fr/jsousa/eVIVALDI. The simulations can be run on a typical desktop computer. In a 3GHz 8-core Mac Pro, with 32GB of RAM, a replicate of a simulation (100 iterations), takes from ~5 to 30 minutes, depending on the parameters. Computations can also be performed in a cluster, allowing the parallel simulation of multiple parameters.

### Entities and their ecological setup

Both bacterial cells and phage particles are represented as independent individuals on an environment represented as a two-dimensional grid with Moore neighbourhood (the 8 connected grid spaces of each location, for a Moore distance of one) (Fig 1A). Bacteria can be of different species. Each individual bacterium has a genome with core, accessory and, eventually, prophage genes. Bacteria have individual phenotypes, such as growth rate or the ability to survive antibiotic exposure. Phages can be from different species, have different lifestyles (temperate, virulent or defective), and possess individual phenotypes (e.g., attachment receptors and burst sizes). The host range of phage hosts is defined by a matrix (Fig 1B), and the superinfection exclusion rules amongst phages is defined in a similar way (Fig 1C).

**FIG 1.**
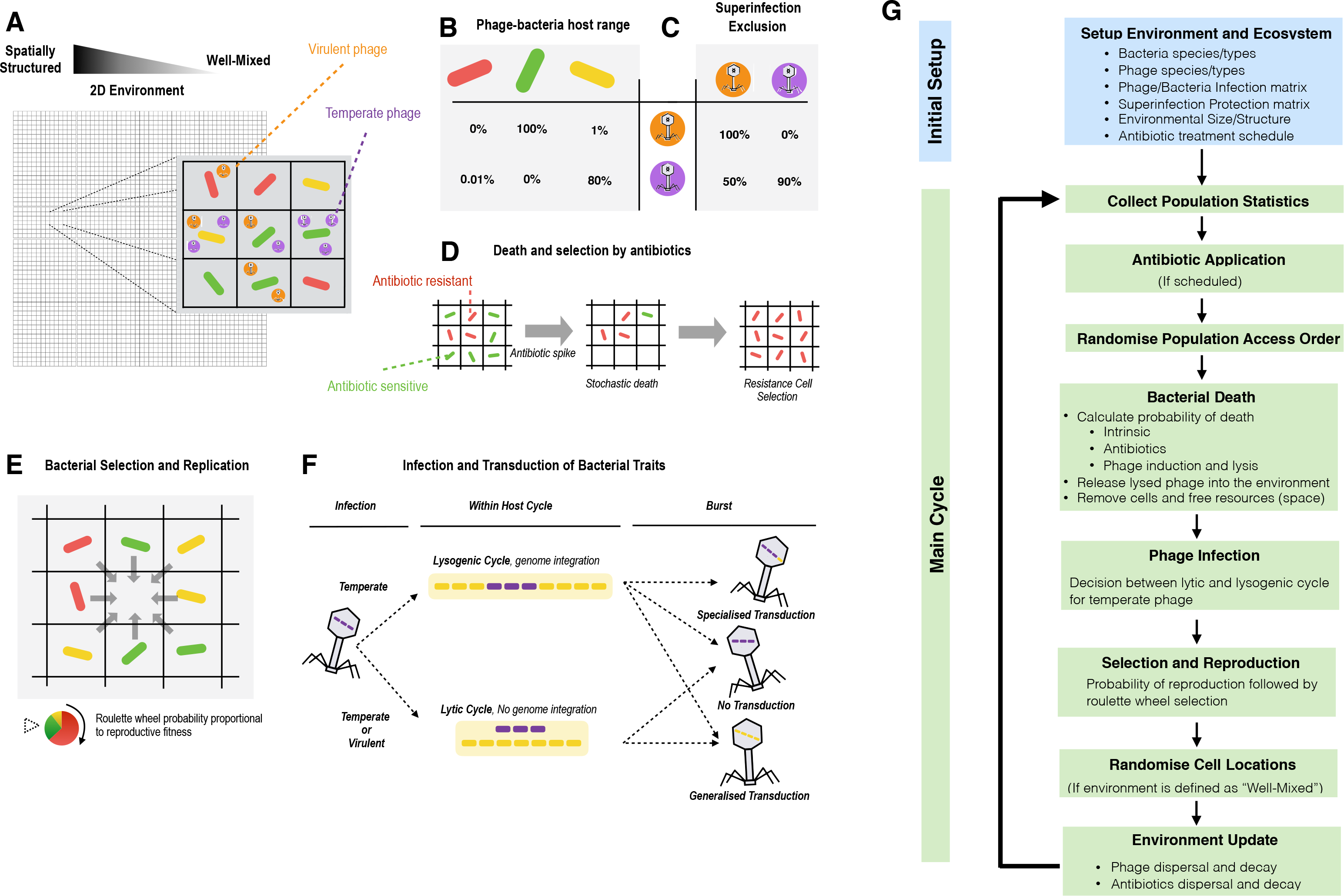
Mechanisms and workflow of the eVIVALDI model. A) Bacterial cells and bacteriophage particles are modelled in a 2-dimensional space, where each (x,y) location holds at most a single bacterial cell and at most a predefined maximum number of phages. The environment ranges from completely well-mixed (liquid), where the contents of each location are randomized at each iteration, to spatially structured, where they are fixed. An intermediate structure is achieved by allowing replication of bacterial cells into a neighbourhood of a given distance. Bacteria and phage can be of different species, and the latter exist as entities either in the environment, where they can infect new hosts, or within hosts, where they either replicate or integrate into their genomes. B) Phage host range is defined in a matrix where each phage has a probability of infecting a given bacterial species. C) Superinfection exclusion is the probability that infection by a given phage aborts when a given type of prophage is present. D) The basal probability of bacterial death can increase by antibiotic exposure or phage infection. Phages decay in function of the period of time spent outside a bacterial host. E) Bacteria compete to reproduce to empty locations, with the fittest bacteria being more likely to produce an offspring. The offspring inherits the traits of the parent cell, but can undergo mutations and is placed into the free location. F) The type of phage infection is determined by the lifestyle of the phage, with virulent following an obligatory lytic cycle, whilst temperate phage can undertake the lytic or the lysogenic cycle following a stochastic decision affected by the density of phages in the environment. The probability of specialized transduction is computed during excision, leading eventually to the incorporation into the phage DNA of a neighboring gene. Generalized transduction occurs before the burst, and a virion has the probability to incorporate random genes from its host, instead of its own DNA. Transduced genes can be used by the subsequently infected bacterial hosts. G) The main cycle of a typical simulation within the model. See complete ODD in Supplementary Material.

### Environmental and bacterial updates

The environment and the individuals are updated and behave according to biologically inspired rules. The environment can be completely structured, semi-structured or not structured at all (i.e., well-mixed), and it can be set as bounded or have a toroidal space. The type of structure influences the diffusion of the different bacterial cells and environmental particles (phage and antibiotics). Each location can hold a single bacterial cell and several phage cells. Free space is the bacterial resource to be consumed, and it is freed whenever bacteria die. Bacterial death can be intrinsic (e.g., of old age) or explicit (e.g., exposure to antibiotics or predation by phage) (Fig 1D). When a free space is available, the neighboring bacteria compete for reproduction. The outcome of the competition is chosen through a roulette wheel method that accounts for the fitness of each bacterium. The successful bacterium generates an offspring into the free space (Fig 1E). Bacteria can be infected by phage in the environment. The outcome of the infection depends on the phage lifestyle and, for temperate phage, the lysis-lysogeny decision. This decision is stochastic but influenced by the number of surrounding phages. For temperate phage, integration in the host genome means vertical inheritance with host replication, until the phage excises from the genome, according to a probability that can be low but non-null throughout the simulation (stochastic prophage induction) and that can also be influenced by the level of antibiotic stress to which the host is exposed. Phage can transduce bacterial genes to other bacteria by generalized or specialized transduction (depending on phages’ characteristics, Fig 1F).

### Input, output and documentation of the model

The inputs of each simulation are two text files that define the general parameters and also the ecological setup of the environment (types and numbers of bacteria and/or phage, along with their attributes). The statistics collected at different time points are stored in dictionaries and dataframes (using *pandas*), can be tailored to the experimenter’s choice and can be represented visually (using *matplotlib* and *seaborn*) or created as an output file.

### Random Forest Analysis

The Random Forest Analysis is based on simulations performed with the model, covering 3000 random combinations of parameters, with 30 simulated repeats per combination. The output of this cohort of simulations is grouped and resumed in response variables, to which a column with 3000 rows of a random parameter is added (i.e., a choice of a number between 1 and 3). This table is used as input of the randomForest package in R (version 4.6.12), where the *randomForest* function is run with the parameters ntrees set to 10000. The relative importance of each parameter (the percentage increase in minimum squared error, %IncMSE) is assessed using the *importance* function from the same package.

### Bacterial strains, antibiotics and growth conditions

Bacteria strains used in this study are listed in Table S2. Bacteria were grown in Tryptic Soy Broth (TSB), Tryptic Soy Agar (TSA) and TSB+0,04%TSA from Oxiod. When appropriate, the following antibiotics were applied: Erythromycin 10mg/L, Chloramphenicol 10 mg/L, Streptomycin 50 mg/L, 0.5 mg/L of Rifampicin. All antibiotics were purchased from Sigma.

### Co-culture experiment

Cultures of JH944, JH927 and JH930 were incubated in 20ml TSB overnight with shaking (200 rpm) at 37◦C. OD600 was measured and cultures were diluted to final OD600 = 0.01 in 1:1 ratios of JH944+JH930 and JH944+JH927 in TSB media supplemented with 5mM CaCl_2_. After overnight incubation (shaking 200 rpm at 37◦C), the cultures were sonicated (10 pulses, 500msec, 50%) to ensure single bacterial cells when plating. Serially diluted cultures were plated on TSA containing erythromycin (JH944+JH927) and chloramphenicol (JH944+JH930) and incubated overnight at 37◦C. The following day, 100 single colonies from the JH944+JH927 culture plates were streaked on TSA supplemented with streptomycin and rifampicin, TSA supplemented with erythromycin and TSA without addition. For the JH944+JH930 culture, 100 colonies were restreaked on TSA supplemented with streptomycin and rifampicin, TSA supplemented with chloramphenicol and TSA without addition. All plates were incubated overnight at 37◦C.

### Phage induction assay

Bacteria lysogenic for phi11 were detected by a prophage induction assay. Bacterial colonies that were present on TSA supplemented with chloramphenicol or erythromycin but absent on TSA supplemented with streptomycin and rifampicin were selected for the phage induction assay. The bacteria were inoculated in TSB and grown to mid-log phase (OD_600_ = 0.5) with shaking (200 rpm) at 37◦C. Mitomycin C (Sigma) 0,4 mg/L was added to TSB to induce any prophages present in the strain and the cultures were incubated at 37◦C with shaking (200 rpm) overnight. The cultures were centrifuged at 4◦C, 3700 rpm for 10 min to pellet the cells and supernatant sterile filtered (0.22 µm). Phages present in the supernatant were detected by mixing 100µl of the filtered supernatant and 100µl of the indicator strain RN4220 in presence of 15µl 100mM CalCl_2_. After incubation at room temperature for 10 min, top agar (TSB+0,04%TSA) was added and the mixture was poured onto a TSA plate supplemented with 10µM CaCl_2_ and incubated at 37◦C overnight. Phages present in the plates were indicative of the original culture being lysogenized by the phage.

## Results and Discussion

### Ecology of phage-bacteria interactions in the light of antibiotic and phage therapy

Antimicrobial therapies rely on the effectiveness of selective agents to kill sensitive bacteria. Phage therapy involves infection and reproduction of the killing agents, thus extending the ability of standard chemical therapies. We started by investigating if eVIVALDI could reproduce simplified but typical ecological scenarios where sensitive individuals are killed by antibiotics and/or predated by virulent phages, thus promoting the increase in frequency of resistant bacteria. A simple community of two bacterial species, one sensitive and another resistant (either to antibiotic or virulent phage), was simulated in a well-mixed environment, and no new resistant bacterial mutants were allowed to emerge in these simulations. Resistance can be defined as costly, in line with experimental data [43], rendering resistant bacteria less competitive in the absence of selection pressure (Fig S1). However, when either antibiotics (Fig 2A) or phage (Fig 2B) were introduced in the environment, the resistant population rapidly increases to fixation. Predation by phage leads to an initial increase in their numbers, because of the abundance of sensitive bacteria, but also to their subsequent rapid extinction when sensitive hosts become unavailable (Fig 2B). A combined treatment of antibiotics and virulent phages leads to the extinction of both populations because none has the ability to survive both selective pressures (Fig 2C). However, the decrease of the antibiotic sensitive population is slower in the presence of phages because of lower competition from antibiotic resistant cells, which are killed by the phage (Fig S2A).

**Fig 2.**
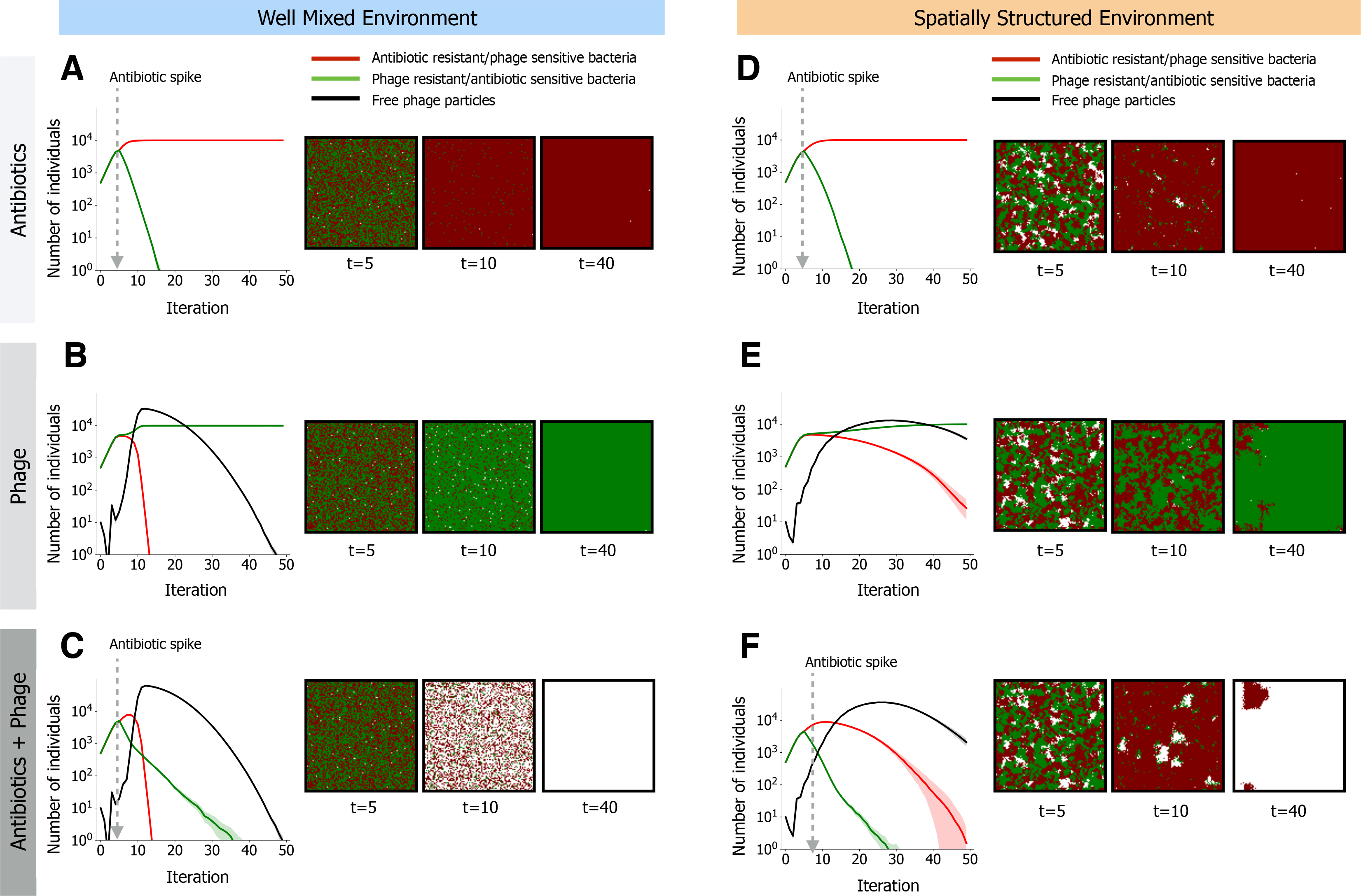
Community dynamics driven by antibiotic selection and phage predation. A small community composed of two different species is subjected to different selective pressures. Bacteria can be sensitive to antibiotics but resistant to phage (in green), or resistant to antibiotics but sensitive to phage (in red). We follow the temporal dynamics and show the populations in their respective colors (the number of free phage in the environment is shown in black). Solid lines indicate mean values for 30 simulations ran with the same parameters and shaded areas show their 95% confidence interval. At the right of each plot is a representative time lapse at 3 time points of the lattices for each scenario, where the colors represent each bacterial species and white spaces represent the absence of bacterial cells. In A) and D) antibiotics are applied at the indicated time. In B) and E), virulent phages (10 individual particles) are co-inoculated with the bacteria at time 0. In C) and F), both selective regimes are applied, with antibiotics applied at the indicated time and virulent phage co-inoculated with bacteria at time 0. In A), B) and C), the environment is homogeneous (well-mixed), as in liquid culture. In D), E) and F), the environment is spatially structured. In D) and F) antibiotics are applied homogenously in the structured environment, and in E) and F) each of the 10 phage particles is initially placed randomly in the biofilm. The complete set of parameters for these simulations is show in supplementary data.

Our model allows to test explicitly the effect of spatial structure on community composition. Spatial structure affects the ability of individuals to diffuse freely in the environment and is known to affect population dynamics [35,36]. Here, spatially structured environments assume that bacteria are fixed in their locations, and can divide only to their immediately adjacent locations. Likewise, we assume that, in these environments, phage cannot diffuse, and thus can only propagate by infecting nearby bacteria. Antibiotics applied homogeneously in spatially structured communities delay the extinction of the sensitive bacteria in comparison to non-structured environments (Fig S2B). However, antibiotics are more likely to be applied non-homogeneously when environments are structured. The delayed extinction is more pronounced in these conditions leading to long term coexistence between sensitive and resistance bacteria (Fig S3). The effect of phage predation on community dynamics is markedly different between well-mixed and spatially structured environments because the latter decreases dispersion leading to “predation waves” that produce spatial arrangements of dead cells akin to those observed in phage plaque assays (see Fig S4 and Video S1). Ultimately, spatial structure results in delayed extinction of phage susceptible cells (Fig 2E vs Fig 2B). Similar to well-mixed environments, presence of antibiotics and phage in spatially structured environments leads to a much slower extinction of antibiotic resistant bacteria compared to environments with antibiotics but lacking phages (Fig 2F vs Fig 2D). However, the presence of phages and antibiotics in spatially structured environments leads to a faster extinction of antibiotic sensitive populations, compared to well mixed environments (Fig 2F vs Fig 2C, Fig S2C), due to a much less efficient phage predation of their competitors when the environment is structured.

The introduction of mutations in the model, eventually reversing the susceptibility to antibiotics or phages, tends to stabilize the bacterial populations (Fig S5A-B). Nevertheless, some populations still go extinct because of the loss of rare mutants by genetic drift or because no adaptive mutations occurred in the time span. Under pressure of antibiotics and phages, double resistant cells emerge only when the mutation rate is very high (Fig S5C). The impact of the environmental structure in the dynamics of predation (Fig 2) led us to analyze how it affects the emergence of resistant lineages (Fig 3). Whilst single resistant mutants increase in frequency slower in structured environments (Fig S5D), double mutants resistant to antibiotics and phage are much more likely to emerge (Fig 3A-D) for intermediate rates of mutation (Fig 3E). This is because in structured environments, the rare mutants resistant to antibiotics benefit from the resources available from neighboring dead cells and rise in frequency without contact with phages (that diffuse less efficiently). This increases the span of time available for the acquisition of secondary mutations conferring resistance to phages, especially if the initial number of phages is not very high (Fig S6). Hence, the acquisition of multiple adaptive mutations is more likely to occur in structured environments.

**Fig 3.**
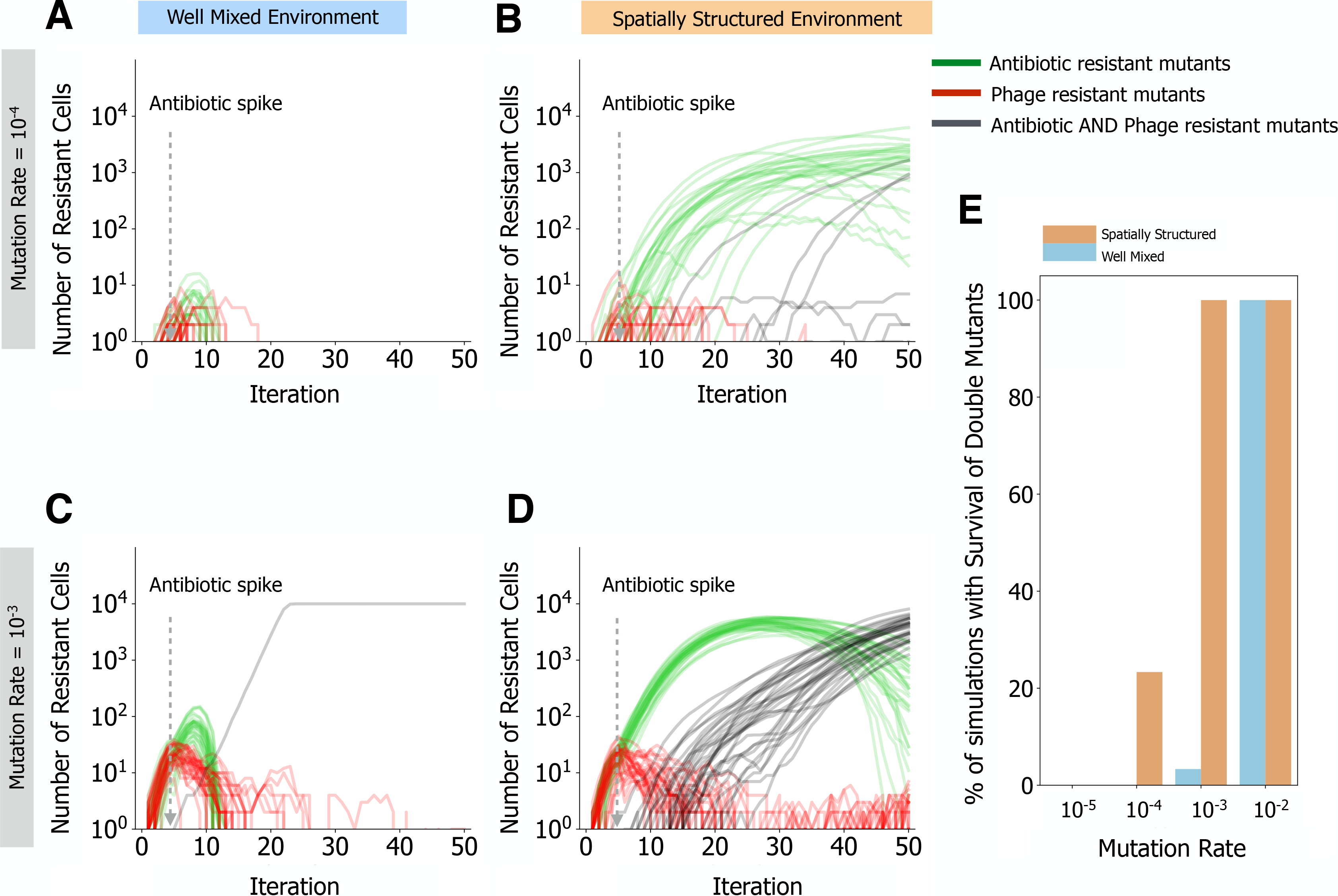
Spatial structure promotes the emergence of multi-resistant bacteria. A-D) Simulations of a single bacterial species, initially sensitive to antibiotics and phage. Lines show 30 replicate simulations with emerging resistant lineages (to one or both selective pressures). Single mutants resistant to phage are shown in red, whilst single mutants resistant to bacteria are shown in green. Double mutant lineages resistant to antibiotics and phage are shown in grey. In A) and B) mutants emerge at a rate of 10^−4^. C and D) mutants emerge at a rate of 10^−3^. A and C show dynamics from well-mixed environments. B and D show dynamics from spatially structured environments. E) Percentage of simulations (out of 30) where lineages resistant both to antibiotics and phage have emerged, in either well mixed or spatially structured environments, for all the mutation rates tested (x-axis). The complete set of parameters for these simulations is show in supplementary data.

The ability of bacteria to evolve resistance to phage might be futile if phage can also adapt sufficiently fast to overcome these changes [44]. When we allowed bacteria *and* phage to evolve in our simulations (Fig S7A), we observed co-evolutionary arms races similar to both theoretical expectations [33] and experimental observations [45]. Spatially structured environments showed slower co-evolution dynamics and higher variability between simulations than well-mixed ones (Fig S7B). Heterogeneous antibiotics added in structured environments further delayed the co-evolution dynamics (Fig S7C-D), due to the death of a significant part of the bacterial population. Importantly, and as before (Fig 3), surviving bacteria (either resistant to antibiotics or not exposed to lethal concentrations) were able to generate mutants resistant to phages for a longer period of time. This is not only due to the limited diffusion of phage, but also because phages need bacterial hosts to replicate and to generate their own genetic diversity. Thus, a reduction in the number of bacterial hosts due to antibiotic exposure hinders both phage propagation and evolution. This suggests that, in natural environments, multiple stressors might render co-evolutionary arms races less predictable than proposed by theoretical models and experimental settings that assume homogeneous populations and environments.

#### Lysogeny as a weapon

Contrary to virulent phages, temperate phages may integrate the bacterial host genome and reproduce vertically with it. The lysis-lysogeny decision in our simulations mimics experimental observations [46], and is influenced by the amount of competition faced by the phage: lysogeny is more likely under high viral concentrations or high multiplicity of infection (Fig S8). Since lysogens are protected from further infections by similar phages, due to superinfection exclusion, the environmental concentration of phages in the simulations decreases rapidly with the increase of lysogens (and depending on free phage half-life). When lysogeny occurs mostly at high viral concentration the bacterial population can become extinct before lysogens can arise. Theoretically, this can also result in the extinction of the phage population.

When a lysogen invader arrives at a community with resident bacteria sensitive to its prophage, lysis of a small fraction of the invaders can dramatically reduce the population of resident sensitive bacteria. This liberates resources for the lysogenic invaders [13]. eVIVALDI recapitulates previous experimental data on this prophage-as-a-weapon hypothesis [17] (Fig 4a): prophage induction rapidly decreases the sensitive population of residents in the early stages of the process, but lysogenization of the latter rapidly neutralizes this process (because the resident lysogens are now resistant to the phage). Hence, the use of prophages as a biological weapon can provide a decisive advantage for colonizing a new niche, but is rapidly neutralized by lysogenization of competitor bacteria. This is also in agreement with previous theoretical works exploring dynamics of invasion in well-mixed environments, using prophages as a competitive weapon [47].

**Fig 4.**
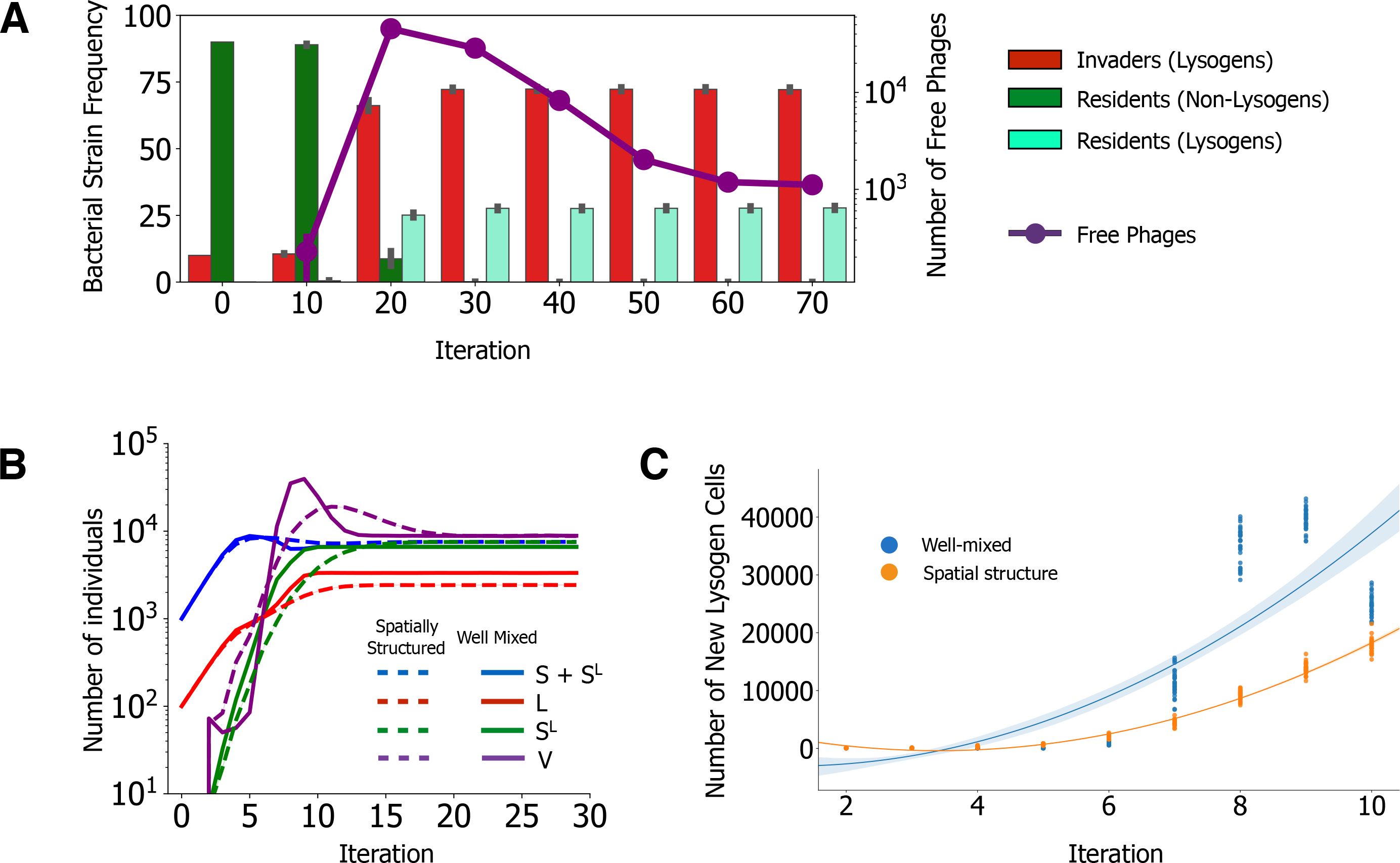
The role of lysogeny in community dynamics. A) Genomes from species A (invaders, grey bars) carry an inducible prophage, whereas those of species B (residents, white bars) are initially non-lysogens. Species are co-inoculated at a 1:10 mixture. Phages (black lines) are spontaneously induced from the lysogenic population. These phages infect the sensitive resident population, which may form lysogens that are protected from phages (B*, black bars). Eventually, the resident that are not lysogens become extinct. All bars represent the average of 30 replicate simulations with similar parameters, with the error bars indicating their 95% confidence interval. Data was displayed as in Figure 1 of [17] for comparison. B) Invading lysogens (L, red lines) and resident sensitive cells (S, blue lines) are co-inoculated at a ratio of 1:10. Phages (purple lines) are spontaneously induced and generate new lysogens in the sensitive resident cells (S^L^, green lines). Full lines: well-mixed environments. Dashed lines: spatially structured environment. Data was displayed as in Figure 3 of [29] for comparison. C) Emergence of resident lysogens in well-mixed (blue) and in spatially structured (orange) environments during the initial 10 iterations of the simulations shown in B. Shown is the polynomial fit of order 2 for the initial 10 iterations, for each of the two types of environment; ANCOVA between the two environments, F=485.5, p=0. The complete set of parameters for these simulations is show in supplementary data.

The advantage of lysogens in the colonization of an environment of resident sensitive bacteria was recently demonstrated in the mouse gut and was suggested to depend on the initial ratio between invaders and resident cells [29]. Indeed, our simulations considering different initial ratios of invading lysogens versus resident non-lysogens showed that the latter were more likely to survive as lysogens when more abundant in the beginning of the process (Fig S9). The abovementioned study presented a population-based mathematical model that fitted well most experimental data, but predicted faster initial infection rates than the observed ones. While different parameters can slow down these dynamics (e.g., the burst size of the phage [29]), the spatially structured mouse gastrointestinal tract is likely to interfere with the temporal dynamics of lysogeny. Interestingly, the inclusion of spatial structure in our model, absent from the abovementioned models, led to a slower increase of free viral particles and slower generation of lysogens in the resident strain (Fig 4B-C). This implicates that invading lysogens may be more successful *in vivo* than would have been predicted by *in vitro* studies in well-mixed environments.

### Autotransduction of an antibiotic resistance gene

When the phages lysing the resident sensitive cells are capable of generalized transduction, they can transfer adaptive traits back to the invader lysogens (autotransduction [27], Fig 5A). To study this process, we started by demonstrating the adaptive effect of lysogenic conversion in bacteria and how it can impact the competition between different phages (Fig S10). We recreated the conditions for autotransduction within our model, by introducing two strains with similar initial population sizes: a non-lysogenic strain resistant to antibiotics ("residents") and a strain of lysogenic antibiotic sensitive “invaders” (Fig 5A). After initial growth, antibiotics are applied in the environment and, as in the experimental study [27], the invaders survive because they acquire the resistance gene by generalized transduction (Fig 5A-B). The analysis of the bacterial genomes in the simulations indicates multiple successive transduction events from the resident to the invader cells (Fig 5C). These events are random (i.e., transduction can transfer any part of bacterial DNA), but natural selection results in over-representation of those transferring antibiotic resistance genes. Overall, invaders lead residents to extinction in most simulations (62%), but sometimes residents become lysogens and outcompete invaders (3%). Interestingly, many simulations exhibited coexistence of lysogenic invaders and lysogenic residents (22%, Fig 5D), and a few showed extinction of all bacterial populations (13%).

**Fig 5.**
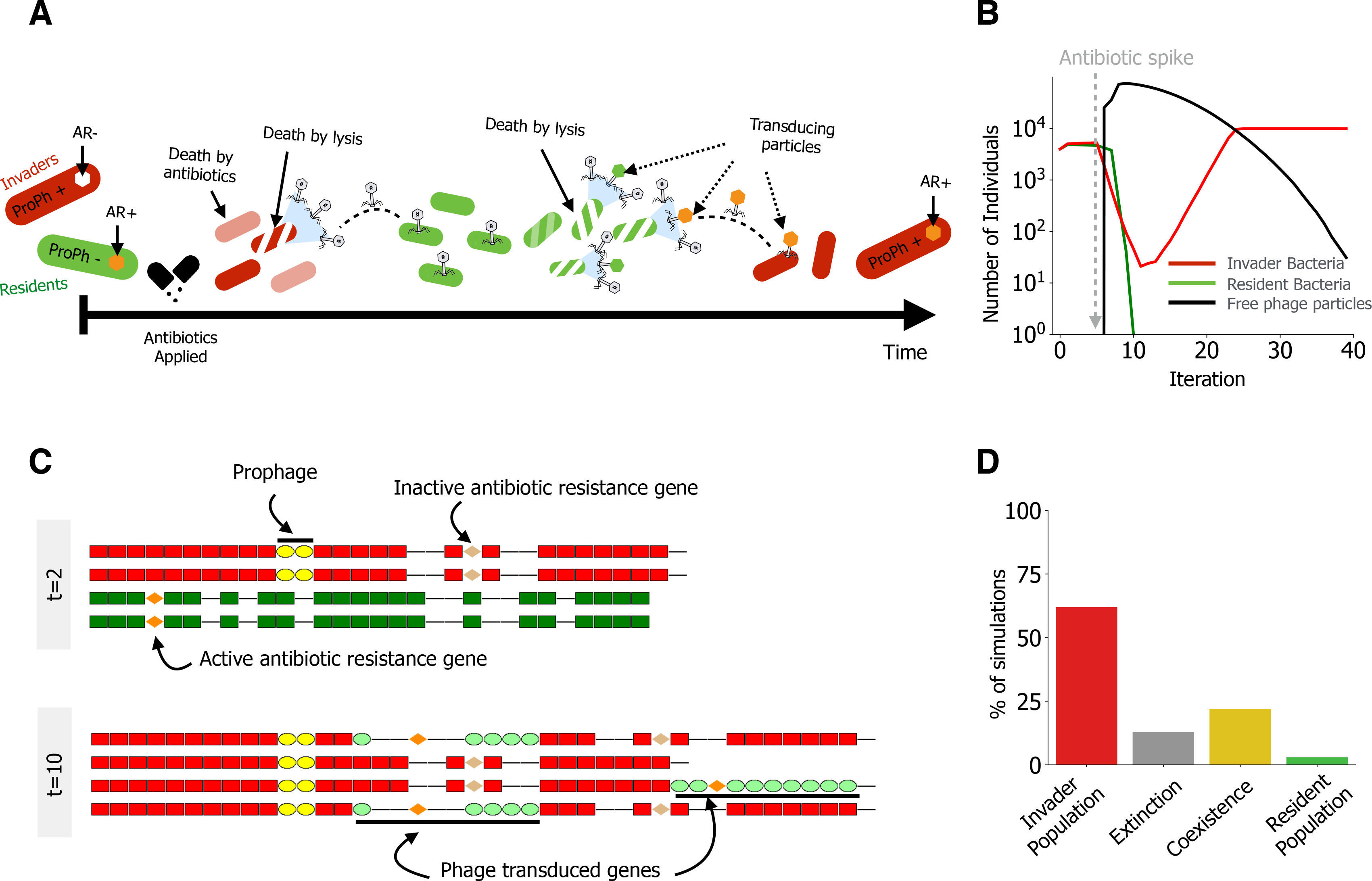
Simulation of autotransduction. A) Representation of the autotransduction events. We created a multispecies community akin to the experimental work of [27], where the invader lysogenic species (red) is sensitive to antibiotics and the resident non-lysogenic species (green) is resistant to antibiotics but sensitive to the phage of the invaders. B) Temporal dynamics of a typical simulation leading to the survival of the invaders. The black line indicates the number of phages in the environment and the time of application of antibiotics is indicated with the grey line. C) Samples of genomes in the population at two different time points of the simulations of panel B. Before antibiotics (*t=2*), the genomes of the resident population (green) carry the resistance trait (orange marker). The invader population (red) is not resistant (grey marker indicates sensitivity to drugs). After the application of antibiotic (*t=10*) most of the invaders have the original prophage and a random sequence of bacterial DNA transduced from the resident cells (other ellipses). D) Outcome of 100 simulations. The complete set of parameters for these simulations is show in supplementary data.

eVIVALDI includes many complex stochastic mechanisms and it is not straightforward to empirically disentangle the importance of each in the final outcome. Therefore, we used a machine learning approach, Random Forest Analysis (RFA, see Methods), to quantify the importance of the mechanisms driving the increase of the population of invaders (Fig 6A, in File S7 are the parameters explored with the RFA). We focused on the percentage increase in minimum squared error (MSE) associated with each variable in the simulation. Generalized transduction had the strongest effect in the efficiency of autotransduction (86% increase in mean square error [MSE], Fig 6B), whilst specialized transduction was almost negligible (3% increase in MSE). Autotransduction also improved with higher probability of adsorption (44% increase in MSE) and infection distance (i.e., the maximum distance between a bacterium and a phage still allowing infection, 70% increase in MSE), because they increase the reach and efficiency of infection by phage and, subsequently, the likelihood of generalized transduction (Fig 6C, Fig S11B). In contrast, when the decision to enter lysogeny (49% increase in MSE) can be made with high probability for relatively low viral concentrations, the resident population proliferates (Fig S11D and Fig S11I for the lysis-lysogeny decision functions explored with the RFA). The importance of the remaining parameters is detailed in Fig S11.

**Fig 6.**
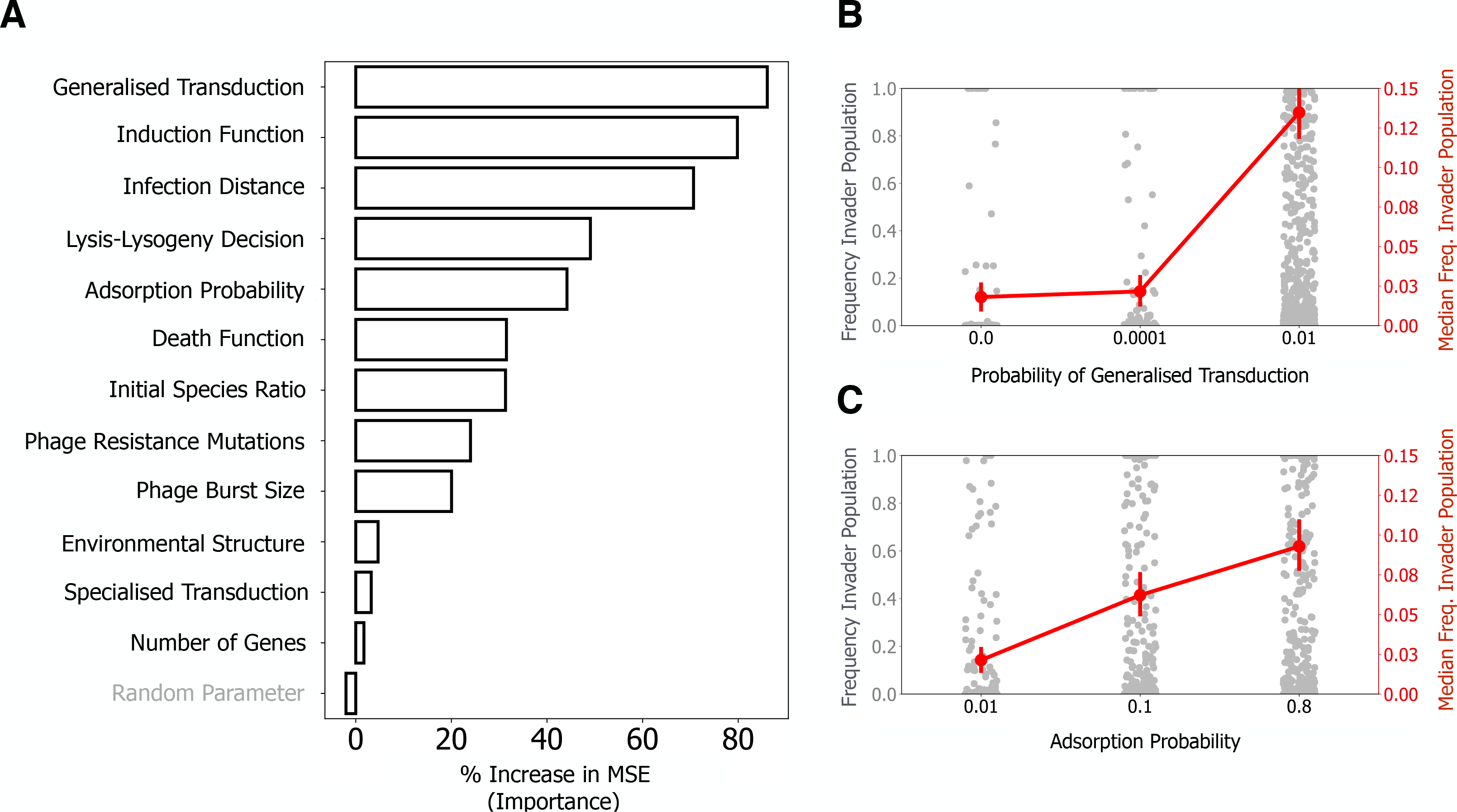
Identification of the main mechanisms affecting the rate of autotransduction of an antibiotic resistance gene using Random Forest Analysis. A) Analysis is based on 3000 randomized combinations of parameters and 30 repeated simulations for each combination). Parameters with a higher % in increased minimum square error have a higher importance for the measured outcome: the median of the final relative frequency of the invader. A random parameter (in grey) was included in the analysis to provide a baseline reference of importance. B-C) The directionality of the impact of two parameters is assessed by plotting the frequency of the invader population at the end of the simulation (across all simulations), in function of the parameter of interest (the other parameters are shown in Fig S11). In the left y-axis, and as strip plot of grey dots, is the distribution of the frequency of the invader population in all simulations. In the right y-axis, and as red dots and lines, is the median of this frequency across the simulations.

To better understand the relationship between two of the most important parameters, generalized transduction and probability of phage attachment, we explored their space of parameters at a higher resolution than before (Fig 7A), while fixing all other parameters. We found that a critical combination of high adsorption efficiency (>1%) and high (between 0.01% and 80%) probability of generalized transduction is required for the survival of invaders (red region). The survival of the resident population is the most likely outcome when rates of transduction and/or infection are low, but also when all phages engage in generalized transduction (100% probability of generalized transduction), because in this case no viable particles are released in the environment (green region). The space of parameters leading to coexistence (yellow region) separates the region leading to the overrepresentation of the invaders from the one leading to the overrepresentation of the resident species (see also Fig S12).

**Fig 7.**
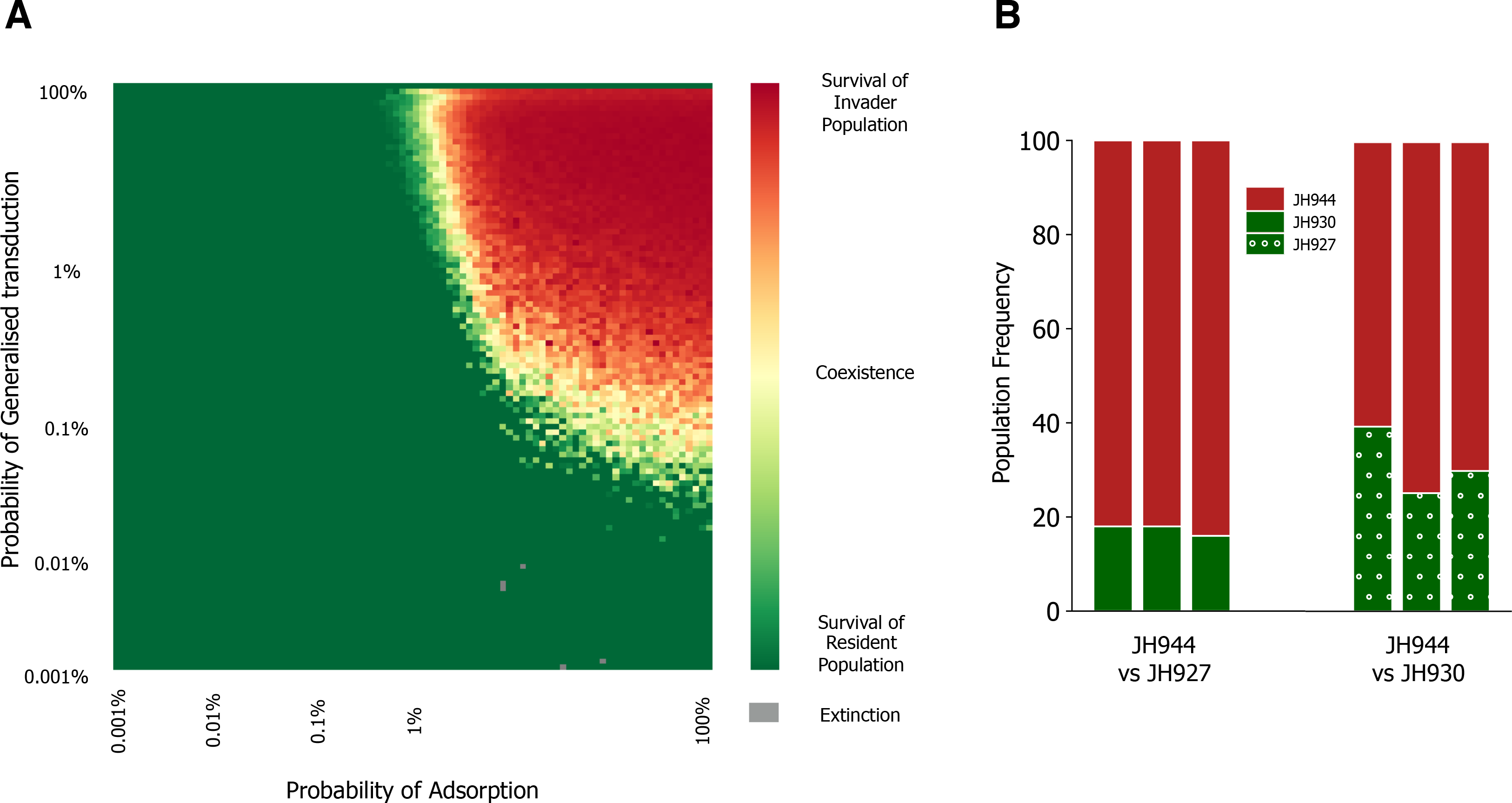
The combined role of probability of adsorption and generalized transduction for the autotransduction of an antibiotic resistance gene. The simulation scenario is similar to Fig 6. **A)** The heatmap represents the likelihood of the outcome of the simulations in function of the two parameters. The color scale ranges from green (100% of the final population composed resident bacteria) to red (100% of the final population composed of invader bacteria), with yellow regions indicating cases where coexistence is the outcome more likely to occur in the timeframe of the simulations. 30 repeat simulations were performed for each combination of parameters, and their median value is used to construct the heatmap. When both populations went extinct, this was either ignored to compute the median (if it occurred in less than 50% of the cases), or was marked as grey (otherwise). **B)** Co-cultures of a lysogenic chloramphenicol sensitive strain of *S. aureus* (JH944, “invaders”, red) and a non-lysogenic chloramphenicol resistant strain (JH930 or JH927, “residents”, green or green with white circles, respectively) indicate coexistence between the two strains at the end of the experiment. Y-axis shows the percentage of colonies with a given genotype (out of 100 or 127 in total, for co-cultures with JH930 or JH927, respectively). Each stacked bar represents an independent replicate of the experiment.

The study describing the discovery of autotransduction focused on the process of gene acquisition by the invaders and did not address the possibility of co-existence [27]. We thus experimentally addressed the prediction of co-existence by co-culturing in liquid media supplemented with chloramphenicol two strains of *Staphylococcus aureus*: a lysogenic strain (JH944, “invaders”) sensitive to the antibiotic and either one of two non-lysogenic strains (JH930 or JH927, “residents”) resistant to the antibiotic (see Methods). The two different resident strains were chosen to demonstrate the result both in the laboratory strain 8325-4 (JH930) as well as in the clinically more relevant strain USA300 (JH927) background. The majority of the colonies resistant to chloramphenicol isolated at the end of the two experiments are in the JH944 background, indicating the acquisition of the resistance gene from the resident bacteria by autotransduction. However, in both combination of strains, performed in 3 biological replicates, a subpopulation of JH927 (17.31.1%) or JH930 (31.36.7%), respectively, were observed to coexist with the invader strain (Fig 7B, Table S3), confirming our predictions. Interestingly, in simulations were coexistence is frequently observed, the frequency of invaders can be influenced by both the probability of adsorption and the probability of generalized transduction (Fig S13). The different frequencies of invaders observed in co-culture with either JH927 or JH930 could represent different regions of the predicted parameter space, where released phages have, for instance, a higher infection efficiency towards JH927 compared to JH930. Confirmation of this hypothesis will require further work on the biology of these phages. Importantly, we observed in the simulations that coexistence is strictly dependent on the generation of lysogenic variants of the resident bacteria, being suppressed when we performed simulations without the generation of new lysogens (Fig S14A-D). This is experimentally corroborated with the observed release of phage particles from the surviving resident clones at the end of the co-culture, when these are exposed to mitomycin C (Fig 7B, Table S3, see Methods). This confirms that lysogenization of the resident bacteria is the mechanism responsible for the coexistence between the two strains and validates the predictions of the model.

Our simulations further suggest that structured environments (Fig S12A) provide an additional region, for extremely high rates of transduction, where coexistence is prevalent (upper regions in Fig S12B-C). These high rates are biologically implausible for viable phages, but not for defective phages or for gene transfer agents [48]. Finally, extinctions of both strains were more frequent when the probability of adsorption was high and transduction was low, suggesting that an inducible phage that is highly infective but a poor transducer is more likely to lead to the collapse of both the invaders and the antibiotic resistant populations. The likelihood of double extinctions is higher in structured environments (Fig S12A, Fig S15 and Fig S16) or in the absence of lysogenization of the resident bacteria (Fig S14E-F and Fig S17). Our results suggest that ecological interactions between strains invading communities of susceptible bacteria can be very diverse, depending on the rates of infection, transduction, lysogenization and population structure.

## Conclusion

Individual-based modelling is providing novel ways to analyze and predict the behavior of microbial systems [40]. Our novel approach integrates multiple and different bacterial species, phages, environmental structures and ecological conditions to explore different aspects of bacteria-phage interactions: temporal changes in community composition (e.g., between lysogens and non-lysogenic bacteria), the concurrent effects of mechanisms of infection, lysogeny, and transduction, and their consequences for the genomic composition of each individual bacteria and phage. To the best of our knowledge, no other theoretical or computation model integrates these different scales of phage-bacteria interactions. This has allowed us to characterize and quantify key ecological components, such as structured environments, in the dynamics emerging from these interactions.

Models are based on simplifying assumptions to make biological systems more tractable. This facilitates pinpointing the relevance of certain mechanisms or agents, but may result in misleading over-simplifications of the system. One major difference between biological systems and our model concerns the number of cells which, due to computational reasons, is lower than the one typically used in experimental settings. Even though our results are qualitatively similar to experimental and/or other theoretical works, this difference may affect the quantitative results. The decreased effective population size (and the consequent increase in the effect of drift) requires that certain rates (e.g., mutation or transduction rates) are simulated at higher values, in order to increase the probability of detecting such events. Another limitation lies in the characterization of the environment. Even if we allow for different levels of structure, environments are spatially and temporally constant throughout the simulations, which might not always be the case in nature. This can change the dynamics of propagation of phages, and lead to subpopulations specialized for different spatial niches. A third limitation of the model lies on the lack of a true physiological description of the bacteria. We assume that phages can infect bacteria at any time, but phage infectivity is known, in some cases, to depend on whether its bacterial host is in exponential or stationary phase [49]. In other cases, a stochastic or induced persistence state in bacteria allows the population to maintain alive a sensitive subpopulation [50]. This can lead to a slowdown or complete halt of infection, particularly in structured environments. Nevertheless, it is important to underline that the model was designed to be easily extensible and further assimilate new mechanisms. Some that are already implemented but not thoroughly explored here include phage resistance based on adaptive immunity (e.g., CRISPR-Cas [32,51]) or mutations affecting the phage host range [52].

One of the major conclusions of this work is that spatial structure affects the dynamics of bacterial populations in the face of antibiotic exposure, phage predation or a combination of both. Whilst combining phages and antibiotics is one of the proposed strategies for the clinical use of phage [53,54], we show here that the emergence of bacteria resistant to both stressors can be enhanced by structured environments, particularly when antibiotics are not homogeneously distributed, as seems common in natural settings [55,56]. This should be taken into account in phage therapy studies.

Adaptation of bacterial cells can also be driven by temperate phage. We showed how autotransduction promotes the spread of antibiotic resistance and is affected by different mechanisms. Importantly, we predicted that different community outcomes (coexistence and extinction) can occur by modulating the efficiency of phages’ infection, lysogeny and transduction, as well as the structure of the environment. We experimentally confirmed the emergence of co-existence between strains of *S. aureus* in well mixed environments, underlining the power of our approach in generating valid and testable hypotheses. It will be important to further test its predictions, as well to simulate other ecologically (and clinically) relevant scenarios. In particular, it will be crucial to explore how the co-existence between virulent phages and prophages influences the outcomes of a combined treatment with phage and antibiotics under a range of ecological interactions between the two types of phages. Exploring these and other ecological settings is also key to understand which factors impact the evolutionary consequences of phage-bacteria interactions for microbial populations.

## ACKNOWLEDGEMENTS

We thank Marie Touchon and José Pénadés for comments, suggestions, and criticisms during the development and testing of the model. This work was funded by European Research Council grant [EVOMOBILOME n°281605] to E.P.C.R. and by an EU FP7 PRESTIGE grant [PRESTIGE-2017-1-0012] from Campus France to J.A.M.S. H.I. is supported by the Danish National Research Foundation’s Centre of Excellence Bacterial Stress Response and Persistence (DNRF12) and J.H. is supported by a Young Elite Researcher grant from the Danish Research Council, Sapere Aude.

